# Assessment of a weak mode of bacterial adhesion by applying an electric field

**DOI:** 10.1101/2020.04.01.020255

**Authors:** George Araujo, Joy Zheng, Jae Jong Oh, Jay X. Tang

## Abstract

Microbial attachment to surfaces is ubiquitous in nature. Most species of bacteria attach and adhere to surfaces via special appendages such as pili and fimbriae, the roles of which have been extensively studied. Here we report an experiment on pilus-less mutants of *Caulobacter crescentus* weakly attached to a plastic surface and subjected to an electric field parallel to the surface. We find that some individual cells transiently but repeatedly adhere to the surface in a stick-slip fashion in the presence of an electric field. Even while transiently detached, these bacteria move significantly slower than the unattached ones in the same field of view undergoing electrophoretic motion. We refer this behavior of repeated and transient attachment as “quasi-attachment”. The speed of the quasi-attached bacteria exhibits large variations, frequently dropping close to zero for short intervals of time. This study suggests applying electric field as a useful method to investigate bacteria-surface interaction, which is significant in broader contexts such as infection and environmental control.

**Significance:** Interaction between bacteria and surfaces occur widely in nature, including those in industrial, environmental, and medical settings. It is therefore important to understand various mechanisms and factors that affect numerous forms of bacterium-surface interaction, particularly those resulting in adhesion or attachment, be they strong or weak, permanent or transient. This work takes a unique approach to identify a transient and reversible mode of bacterial attachment to a solid surface, by applying an electric field to exert a force for detachment. The force thus exerted proves to reach the amplitude required to detach bacteria of a pilus-less strain that weakly attach to a plastic surface. The method may be applied broadly to investigate bacteria-surface interaction.

## Introduction

Ubiquitous in nature, microbial attachment to surfaces has been widely studied^1–3^. To human and animal hosts, attachment is often the first step towards bacterial infection^4^. Bacterial attachment is also the most crucial step for the formation of biofilms^5, 6^, which may also involve other triggers such as fluid flow^7^. In the process of exploring their environments, bacteria often encounter a boundary. In spite of swimming motility and Brownian motion^8^, motile bacteria tend to accumulate near a boundary surface^9–11^. When within 2-3 *μ*m from a solid surface, they may become sterically or hydrodynamically trapped and remain in the proximity of the surface for significant lengths of time^9, 12^. Whereas near surface accumulation and entrapment help facilitating bacterial adhesion, these interactions are distinct from the actual attachment.

Attachment to surfaces can be classified according to its molecular basis, as well as strength and reversibility. Most bacterial species attach and adhere to surfaces via special appendages such as pili or fimbriae. The initial attachment by pili and their retraction are known to lead to strong and permanent attachment, the mechanism of which has been extensively studied^1^. Less is known, however, for other forms of attachment, particularly those weaker and more reversible, even though they play functional roles. For instance, weak and reversible adhesion has been shown to enhance surface colonization^13^. Sequences of attachment and detachment of binding tethers may serve as pre-play in the process of consolidating irreversible attachment^14^. These transient attachments are also linked to optimal surface exploration as it has been proposed to maximize bacterial surface diffusivity^15^.

Bacterial colonization of a surface can be undesirable, such as in formation of dental plaques and various types of bacterial infection^16^. As a consequence, different techniques have been applied in order to remove bacteria from surfaces. The most common examples involve flow, which may be produced by peristaltic pumping or through passage of air-bubbles^17^. Bacteria may be dislodged from conductive surfaces by applying an electric field^18, 19^. It was observed in another study that electric forces parallel to the surfaces are more efficient to promote detachment than those applied in the perpendicular direction^20^. However, there has not been much study to quantitatively assess bacterial attachment to surfaces by application of electric field.

We chose a Gram-negative bacterium called *Caulobacter crescentus* as a model system for the study of bacterial adhesion. Commonly found in aquatic medium, including soil and drinking water^21^, *C. crescentus* has a dimorphic life cycle: One newly divided cell is motile and called a swarmer, which in about 45 minutes sheds its single flagellum, hence losing its motility. It then develops into a stalked cell. Under favorable conditions, the stalked cell proceeds to the next division, producing a motile swarmer cell as its offspring in every two and half hours^21, 22^. The stalked cell is known to produce a holdfast capable of strongly and permanently adhering to a solid surface^23^. However, in most laboratory based adhesion studies of *C. crescentus*, the initial attachment tends to occur early in the swarmer stage. Effective attachment to surfaces for this species is aided by the presence of type IV pili and flagellar motility^24^. In fact, the swarmer attachment has been shown to trigger just in time expression of holdfast and accelerate the swarmer cell to stalked cell transition^25^. Aside from strong adhesion and permanent attachment via its holdfast, facilitated by pili, we focus in this work on mutants that lack pili in order to study weak attachments involving other surface molecules or mechanisms. A recent study on the tethered motion of *C. crescentus* swarmer cells of the same strain suggests a “fluid joint” possibly formed by polymers on the cell surface^26^. In addition, it has been reported that the curved shape of *C. crescentus* may help the cell to colonize surfaces^27^.

In this study, by applying an electric field parallel to a plastic surface, we perform experiments based on the initial observation that some weakly attached cells are readily detached while others remain stuck to the surface. More interestingly, a small number of tethered cells exhibit the characteristics of moving slowly along the surface, driven by the field, but never leaving the surface proximity. These particular cells move at erratic speeds. They alternate between being transiently mobile and tethered in a stick-slip fashion. We refer to this phenomenon as *quasi-attachment*. This report focuses on this peculiar behavior and explores the mechanism for it.

## Results

### Types of motion under an electric field

Under an electric field, four different behaviors were observed for individual cells: 1. Non-swimmers, including stalked cells and possibly some dead cells, drifted towards the positive pole, driven by the electrophoretic force caused by the electric field. 2. Cells that remained nearly immobile for some time, at most giggling about their fixed positions due to Brownian motion. Some of these cells were seen to be spinning at fixed positions before an electric field was applied, an indication of surface attachment through body or flagellum tethering. 3. Swimmers that kept their swimming motion, either against the direction of the field (more often) or opposite to it. 4. a small number of Cells, which initially attached to a solid surface, moved along the field slowly in a slip and stick fashion. The first three types of motion are more common and better understood, which are briefly assessed in the next sub-section. The type 4 behavior, both surprising and significant, is shown separately and then contrasted by results of a control experiment.

### Alignment of cell trajectories under an electric field

In the absence of external electric field, the swarmer cells of *C. crescentus* swim in random directions, which change stochastically in the time scale of seconds (Figure 1a). Once exposed to an electric field, the effect of galvanotaxis is immediately observed. The applied electric field imparts a drift speed on every bacterium, motile or not, but the effect on swimming cells is more dramatic as the field has aligned their swimming trajectories. The average direction of motion is opposite to the direction of the electric field, indicating negative net charge of the bacterial cell. The alignment in trajectory is mainly caused by orienting the cell-flagellum axis, with the cell body heading towards the positive electrode and the flagellum pushing from behind in most cases. This observation suggests the cell body is more negatively charged than the flagellum, and the ratio of the electrophoretic driving force on the body to that on the tail is larger than that of the hydrodynamic drags on them. Under stronger fields, the paths of motion become closer to long and straight lines that are nearly anti-parallel to the field direction (Figure 1b).

**Figure 1.**
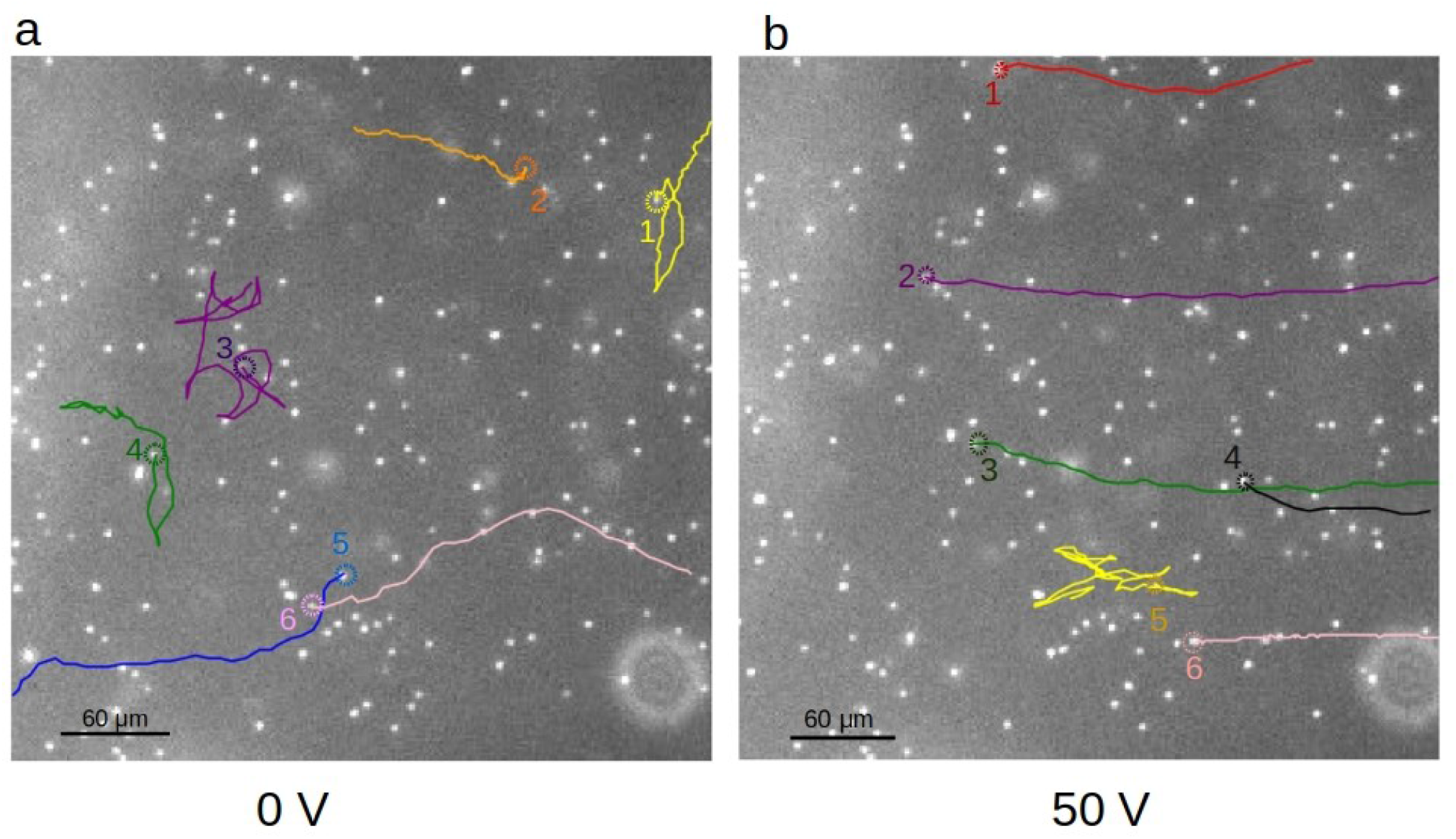
Bacteria trajectories. a) Trajectories of several CB15 ΔPilin swimming cells in the absence of electric field. Note numerous reversals in swimming direction for cells 1, 3 4, due to frequent switches in flagellar motor rotation. b) Under 50 V voltage applied, bacteria move predominantly against the applied field. The direction of the field on the figure goes from right to left. The dashed circle on each trajectory indicates the starting point of the track. The bright dots are images of bacteria, which were either stuck to the surface or moving near the surface. For clarity, only six cells are highlighted on each figure. Some cells moved beyond the borders of the imaging area. The images were taken using a 4x objective lens.

It is interesting to note that some cells are observed to swim in the direction of the field, i.e. towards the negative pole, typically at much slower speeds. In these cases the swimming direction is parallel to the electric field. We know from previous studies that when the flagellar motor of *C. crescentus* rotates counter-clockwise, the flagellum pulls the cell body backward^8, 28^. This backward swimming motion appears to have overcome the electrophoretic effect, resulting in net slow motion down stream with respect to the electric field. In the absence of electric field, *C. crescentus* switches the direction of swimming within a few seconds^29^ (unless genetically modified to prevent the switching behavior, such as the SB 3860 strain, which swims solely forward). We indeed observed switches under the applied field, but an arbitrarily large change in the direction of motion as the result of a flick^30^ became less evident. Even with changes in the direction of motion, the trajectories remain aligned with the electric field. One example is shown as trajectory 5 in Figure 1b. This alignment behavior is not specific to *C. crescentus:* our limited experiments with *Pseudomonas aeruginosa* yielded similar results. In short, immediate and large changes in trajectories of swimming bacteria under an applied electric field can be accounted for by taking into consideration both flagellum propulsion and field alignment due to electrophoretic forces on the bacteria.

### Speed variation of weakly attached cells moving under the field

Unattached cells, whether swimming against the direction of the electric field or solely moving electrophoretically, travel faster as the field strength is increased. In contrast, the slowly dragging cells behave differently. These cells transit from short moments of limited mobility to moments of negligible mobility. We define the weak interaction of these cells with the plastic surface as “quasi-attachment”. Figures 2 a,b highlight an example of this behavior (labelled as cell 2), in contrast to cells that are either permanently attached to the surface (cell 1), or drifting electrophoretically (cell 3). Supplemental movie S1 shows a video from which the images were taken. While cells freed from the surface attachment quickly move away from their initial positions, the motion of a quasi-attached cell is much slower, as indicated on the images, with displacement and speed plotted in Figure 3.

**Figure 2.**
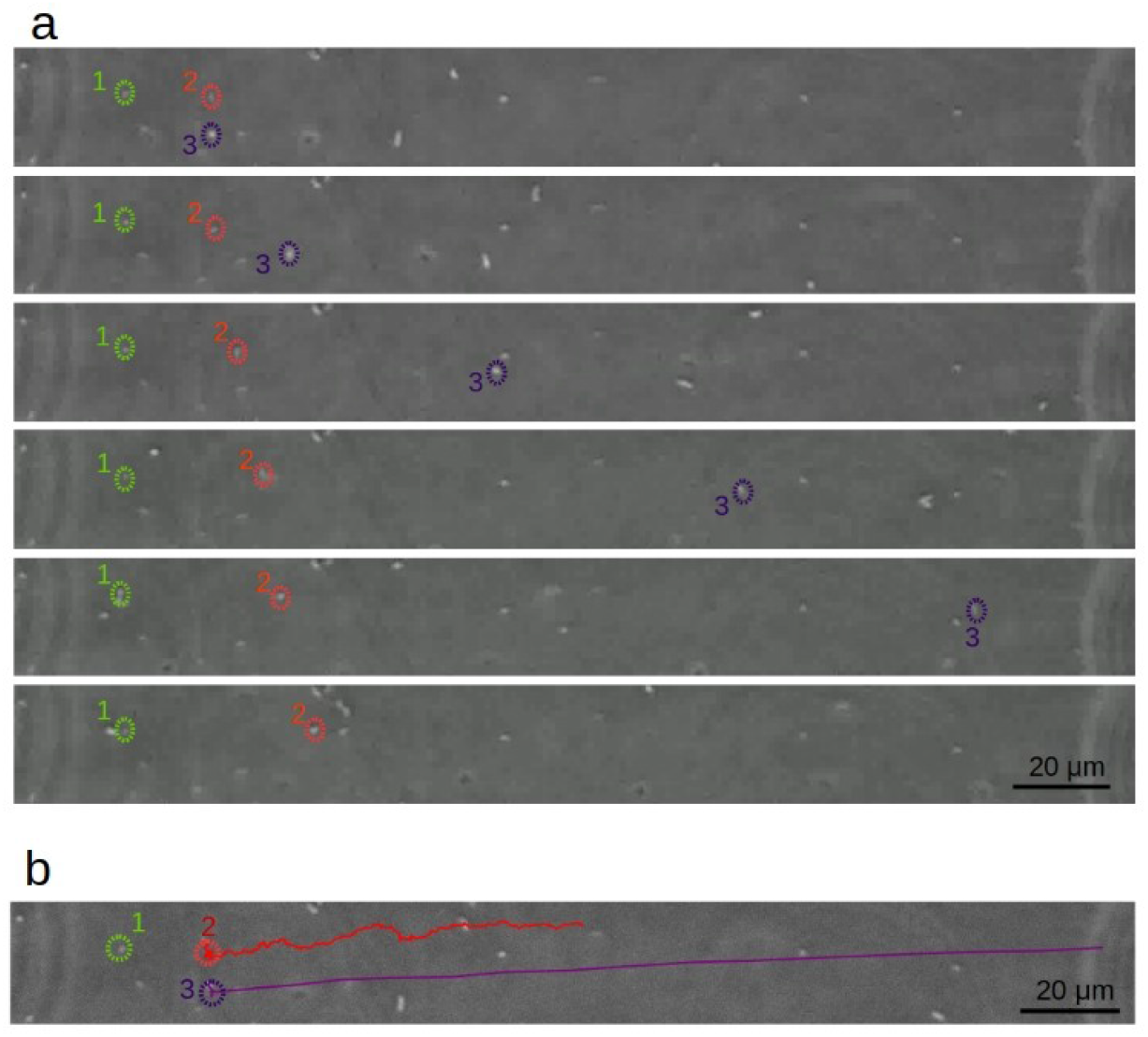
Types of bacterial motion under DC electric field. a) Cell 1 is attached to the surface and hardly moves. Cell 2 moves slowly under the field (quasi-attached), making frequent changes in speed. Cell 3 is initially attached to the surface, but it detaches quickly after the electric field is applied and moves fast and freely along its path. The images in the sequence are 2 seconds apart. b) Tracing the trajectories as illustrated in Figure a) over 31 seconds. Note that the time interval traced is longer than what is shown in the time lapse sequence in a). These images were taken using a 40x objective.

**Figure 3.**
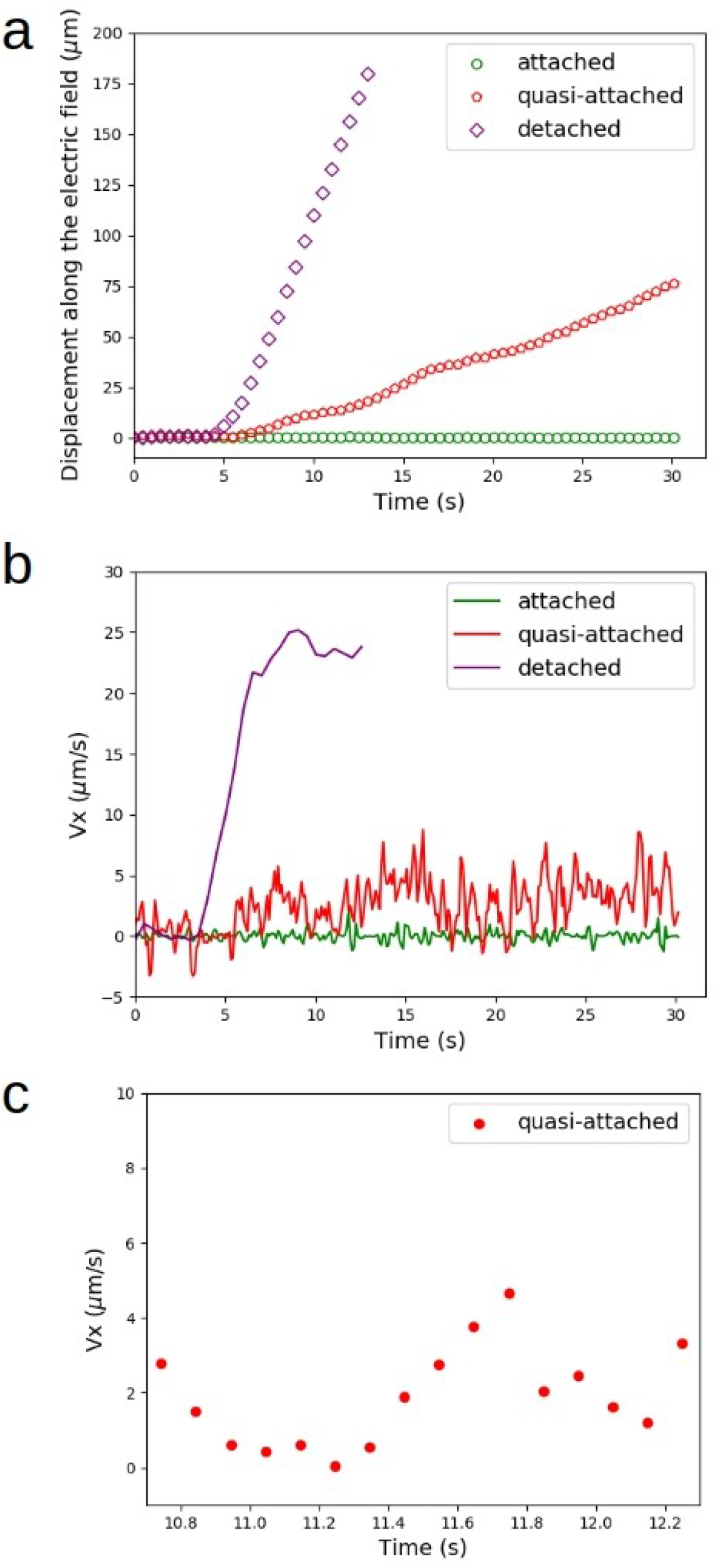
Displacement and velocity along the field. a) Displacement of the highlighted cells along the axis of the electric field. Experimental data are shown for points separated by 0.5 seconds, which is five times the time interval for data acquisition. The voltage applied was increased from 0 V to 100 V within seconds, and then kept at its maximum value. b) Velocity along the field as a function of time for the three highlighted cells shown in Figure 2. c) Zoomed in plot of velocity values lasting approximately 1.5 seconds of the experimental data in Figure *b* above. Within this short time interval, we see in more detail the variation of velocity along the field. At the moment from 10.9-11.3 sec, for instance, the velocity was close to zero, indicating moment of transient attachment.

During the short intervals when a quasi-attached cell was moving, its velocity component along the field axis remains highly variable, as is shown in Figure 3b. It is possible that this noisy speed-time curve was also due to very brief moments of attachment, but our method of observation could not confirm that due to low frame rate. In this study, all speed values were averaged over about one tenth of a second, which was set by both pixelation accuracy and the frame rate of the images taken.

One surprising behavior of these bacteria in draggy, stick and slip type of motion is that no obvious correlation is noted between the intensity of the applied field and the average moving speed of the cells. Figure 4 shows six examples of averaged velocities along the electric field axis from the quasi-attached cells (strains CB15 Δ*Pilin* and SB 3860) under different electric fields. There is a large variation in each cell’s moving velocity over time, as indicated by the large error bars representing the standard deviations. The average velocity also varies among the 6 cells, but there is no obvious correlation between the average velocity and either the voltage applied or the strain of the cells tested. Note the trajectories of these cells are aligned with the field axis. Thus the velocity component along the field axis properly accounts for the average drift velocity of these transiently attached cells.

**Figure 4.**
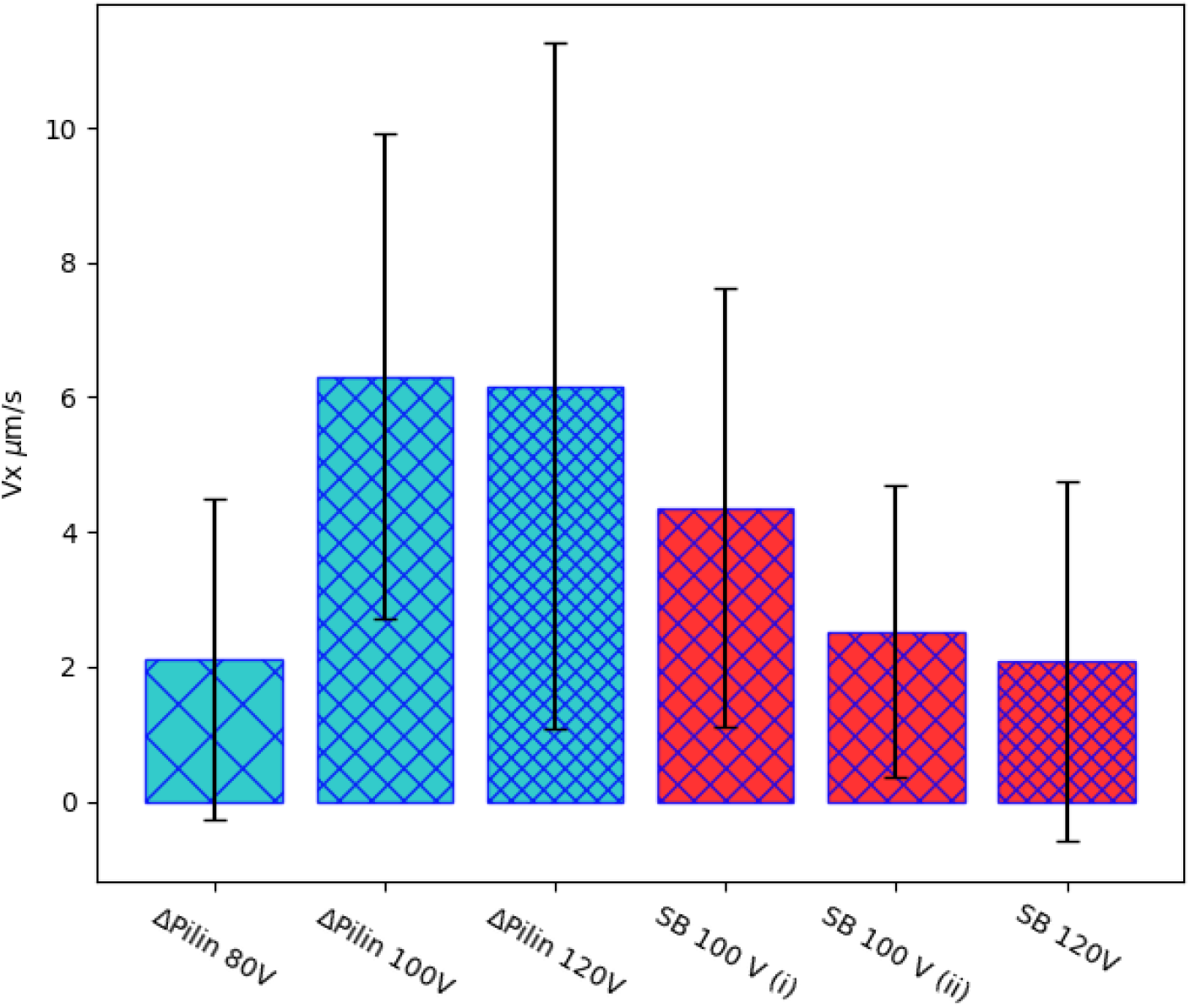
Average velocity component along the axis of the electric field for six dragging cells from two different strains and under three different values of applied voltages. The error bars represent the standard deviations.

### Control experiment of polystyrene beads transiently attached to the same plastic surface under electric field

In order to discern to what extent the quasi-attachment of bacteria on the plastic surface might be caused by non-specific physical forces, such as that attributable to the classic Derjaguin-Landau-Verwey-Overbeek (DLVO) theory^8, 31^, we performed a control experiment using colloidal beads under the same setup. We used polystyrene beads comparable in size to that of bacteria, diluted in the same solution as used for the bacteria. We found that a much lower voltage was required to dislodge the small number of beads that occasionally attached to the plastic surface. The range of voltage was 15-25V, as opposed to 80-120V for the bacteria. Such a difference reflects the different amounts of surface charge carried by these distinct classes of particles. Nevertheless, we found a small number of them in both cases transiently attached to the plastic surface, thereby allowing for a close comparison.

The beads near the plastic surface behave very differently from the bacteria, motile or not. Most beads drift electrophoretically under the applied field. In rare instances, some beads become transiently immobile for variable intervals of time on the order of seconds. After freeing itself from transient attachment, the bead usually resumes its electrophoretic motion of the same velocity as before, and comparable to the majority of beads that never become attached. Figure 5a shows a trajectory of a transiently attached bead in comparison with a freely drifting bead under an applied voltage of 15V. The velocity profiles of these two beads are plotted in 5b. Two other examples are shown in separate experiments under slightly higher voltages applied (5c and d). All these examples show a very different behavior from the quasi-attachment of bacteria. Whereas the latter displays dragged motion of nearly continuous attachment, the transient attachment of a bead is binary: it appears either totally attached or detached, and there is no lingering effect on the bead’s drift speed once it is detached.

**Figure 5.**
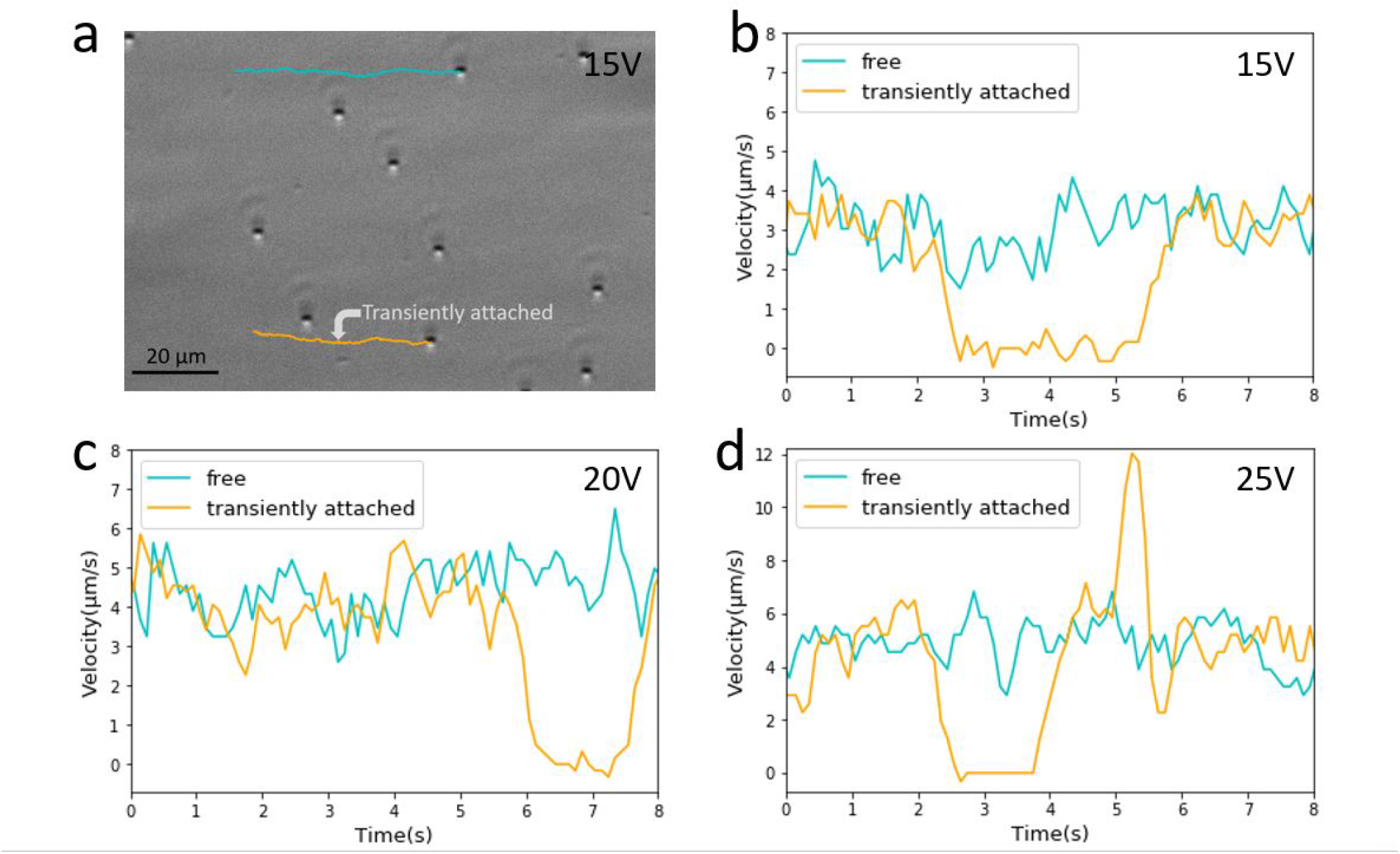
Transient attachment of colloidal beads in a control experiment. a) First frame of a video capturing electrophoretic motion of beads under 15V applied voltage. Superimposed are two trajectories, one of a freely drifting bead (cyan), and the other also drifting (yellow) except for a 3-second pause at a spot indicated by an arrow. b) Velocity versus time plots of two beads from their trajectories indicated in a. c and d. examples of similar beads motion observed in two other experiments under slightly higher applied voltage values of 20V and 25V, respectively. The images were taken using a 40x phase contrast objective lens. The positions of beads were captured in 0.1 sec intervals, but the velocity values were 5-points running averages in order to reduce the noise in display.

## Discussion

### Charge distribution along the bacterial cell is primarily responsible for trajectory alignment

The cause for the alignment of bacterial trajectories under an electric field is physical in nature: there is a difference in electrophoretic mobility of the cell body from that of the flagellar filament^32^. The typical surface area of the cell body is in the range of 2-5 μ*m*^2 33^, which is around two orders of magnitude bigger than the surface area of the flagellum, whose averaged contour length is a few μm but cross-sectional diameter only about 14 nm^34^. The hydrodynamic drag on the cell body is also larger than on the much thinner flagellum. Based on our observation that motile cells swim rapidly towards the positive electrode, faster than the drift of non-motile cells, we know that the cell body usually leads the flagellum. In this orientation, the bacteria are both driven by the field and pushed by the flagellum towards the electrode of higher voltage. The experimental results also suggest that the ratio of the drag coefficient is much less than that of the effective charges between the two. The imposed cell orientation dictates the kinematics of the bacterial trajectories. Under strong fields (with applied voltages above 40V in our experiments), cell trajectories do not deviate much from straight lines even for cases when we observe switches in the direction of motion. This result is due to the alignment of the cell body by the electric field, which may also suppress the flagellar flicks^28^. For *C. crescentus*, with its flagellum of right handed helical structure, a flick may occur as its flagellar motor switches rotation from counter-clockwise to clockwise, causing the cell body to change from backward to forward motion. Whereas a flick often causes the bacterium to reorient its cell body orientation, causing a large change in swimming direction in the absence of applied field, the presence of a strong electric field keeps the cell body aligned, thus suppressing the randomizing changes in swimming direction caused by the flicks regardless whether or not the frequency of switches in motor rotation directions is affected.

### Comparison on forces of detachment by several methods

Applying an electric field to detach cells from a surface offers a unique range of force as compared with other techniques. In our experiment, the typical force applied is in the range of 0.1-0.5 pN (estimated in supplemental document). Such a strength of force is 1-2 orders of magnitude stronger than achieved by a common micro-fluidic approach^35^. However, it is 1 to 2 orders of magnitude smaller than that can be exerted by laser tweezers (1-100 pN)^36, 37^, and several orders of magnitude smaller than forces applied by atomic force microscopy (AFM) (orders of pN-nN)^38^ or micromanipulation techniques (up to *μ*N)^23^. The strength of bacterial adhesion spans such a wide range of magnitudes, which has been demonstrated by experiments applying all these techniques. Interestingly, the present study shows that forces in the sub-piconewton range are required to detach the swarmer cells of two pilus-less strains of *C. crescentus* that weakly and non-specifically attach to a plastic surface. Such a weak interaction might be significant prelude to much stronger and permanent attachment to surfaces mediated by pili and/or other appendages. There has not been that many studies in this particular range of interactions, perhaps due to a lack of appropriate tools. Although relatively weak attachment, what we observed and measured in this study may be among the interactions that are stronger than those that can be conveniently washed off by moderate shear flow, limiting the effectiveness of conventional micro-fluidic devices (See a comparison between the electric force and shear force in Supplemental Information). Instead, the method we have shown in this study may prove useful for future studies on a variety of microbes in the context of their attachment and adhesion to surfaces, including those under environmentally or medically significant settings.

### The relevance of DLVO theory to weak and transient attachment

One previous study explored the role of cell surface lipopolysaccharides in *E. coli* adhesion to the surfaces of quartz beads or coverslips^39^. Using packed bed columns and a radial stagnation point flow setup, the study investigated the transient interaction of K12 strains expressing several lengths of polysaccharrides under different ionic conditions with silanized surfaces. The authors suggest combined influence of electrostatic interactions, in terms of the classic DLVO theory, LPS-associated chemical interactions, and hydrodynamics of the deposition system.

The DLVO energy is given by adding the van der Waals and electrostatic contributions for the bacterium-surface interaction. For a sphere near a flat surface, these energies are given by equations 1 and 2^31^.

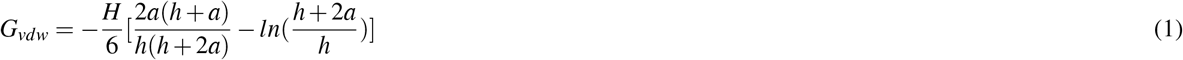

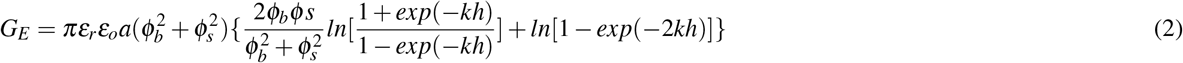

In the equations above, H is the Hamaker constant, h is the bacteria-surface separation, *a* is the bacterium radius, *k* is the inverse of the Debye length, *ε_r_ε_o_* is the dielectric permittivity of the medium and *ϕ_b_* and *ϕ_s_* are the bacterium and the surface potentials, respectively. The derivative of the total energy with respect to the sphere-surface distance (h) yields the interaction force.

Using typical values to represent these quantities^8^ (for glass surface in this reference study), the DLVO force is estimated to be around a fraction of pN when the system is at the vicinity of the second minimum of energy (noted from a plot in the published supplemental materials of the said reference). This estimate coincides with that for the force provided by the electric field, supporting the idea that while dragged along by applied electric field, bacterial cells, like colloidal beads, may become weakly attracted to the solid surface. Aided by incessant Brownian motion, they may occasionally bump into and become transiently attached to the surface.

However, the strikingly different behavior between the transient attachment of colloidal beads and quasi-attachment of the pilus-less bacteria on the same plastic surface points to the necessity of additional, more specific mechanism than DLVO, which applies to both but could not account for their qualitatively different behaviors. Thus, the ensuing discussion examines other hypotheses, followed by a model we propose to account for the stick and slip type bacterial attachment.

### Assessment of Alternative Hypotheses

One simplistic hypothesis is that these slow moving cells are just cells that are swimming towards the negative pole, i.e. in the field direction, opposite to the electrophoretic driving force. In this scenario they would be moving slowly because they were “fighting” against the electric force. However, since the slow irregular motions are seen over dozens of seconds, whereas we know from an earlier study that the characteristic time for switching the direction of motor rotation rarely goes beyond a few seconds^29^ (for Δ*Pilin* cells; SB3860 only swims forward). It would be very unlikely based on this picture that a Δ*Pilin* cell would be swimming over 30 seconds in the same direction along the electric field. In fact, we observed multiple examples of slow, dragged motion for Δ*Pilin* cells, thereby ruling out this model. On a separate line of reasoning, this hypothesis can also be ruled out based on the fact that we observed these slow and dragged motion at several applied voltages. It would be rather hard to achieve the nearly exact cancellation in all these experiments since the electrophoretic velocity is proportional to the applied voltage, whereas swimming speed would remain constant.

A second hypothesis is that the no-slip boundary condition for fluid flow might account for the transient and slow dragged motion, which we noted “quasi-attachment”. In other words, the observed slow motion could just be reflecting the extremely slow-flowing fluid containing non-motile cells very close to the surface. This picture is inconsistent with results from an earlier study measuring the swimming speed of *C. crescentus* as a function of its distance from glass surface using total internal reflection fluorescence (TIRF) microscopy. In that study, the cells were found to swim relatively freely as close as on the order of ten nanometers from the surface, with their typical swimming speeds ~40 *μm/s*^8^. The qualitatively different type of motion and the much slower and highly variable motion of the subset of cells identified in this study lead us to other mechanisms.

Another possibility is that the flagellar filament frequently scratches the solid boundary, causing too large a drag to allow fast motion of those cells touching the surface. Given that a typical flagellar rotation rate is in the range of 250-375 Hertz^40^, the flagellum could be interacting with the surface on the order of hundreds of encounters per second at some rough spots, imposing a strong drag on the bacterium. This interaction of flagellum and surface could also be a cause for the quasi-attachment. A potential method to test this hypothesis would be by directly visualizing flagellum. The method would require labelling the flagellar filament while not severely affecting the flagellated motion. One might then be able to directly visualize the behavior of the labelled bacteria when they display quasi-attachments under applied electric field. This approach has been attempted previously, using genetically modified strains of *C. crescentus* so that its flagellum could be labelled. Unfortunately, the labelling resulted in much reduced motility and only in very few cases, short video segments showing in rare occasions a motile cell with a faintly labelled filament^26^. Once a better labelling technique is developed, applying it to the experiments under electric field as described in this report could help to clarify to what extent the flagellar filament affects the quasi-attachment.

### The proposed mechanism

We propose that cell surface polymers, such as long and flexible polysaccharrides richly expressed on the bacterial surface, account for the quasi-attachment of pilusless bacteria to a solid surface. This attachment is similar to the “weak rolling mode of surface adhesion” for *E. coli*^13^, yet the molecular origin may be different as the latter is known to bind to mannosylated surfaces via the adhesive protein FimH. The molecular mechanism we envision is akin to what has been proposed in a recent report^14^, albeit on different species of bacteria.

It has been proposed^26^ in previous report based on experiments on the same *C. crescentus* strains that we study that a “fluid joint” made up of polysaccharrides that cover the cell wall could be responsible for the interaction with the surface (Figure 6). The tether region seems to be located near the flagellar pole in most cases, but occasionally also near the center of the body^26^. Given that the nature of this cell-surface attraction is proposed to be caused by polymeric link between the bacterial cell wall and the surface, we suggest to refer this interaction as via *polymeric tethers*^39, 41^. The reason for this proposed change in terminology is that “fluid joint” implies no molecular link, but instead, the action of a thin layer of confined fluid. We hereby suggest explicitly in the sketch multiple molecular contacts, which collectively account for what we refer as “quasi-attachment”.

**Figure 6.**
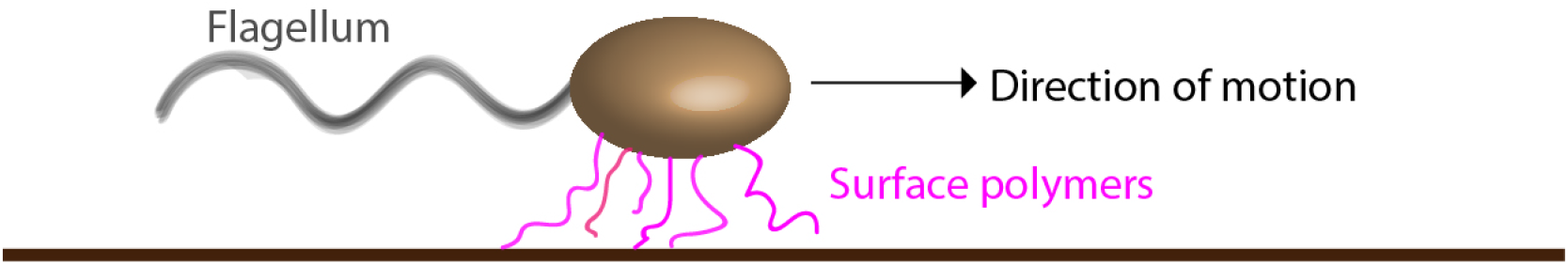
Illustration of quasi-attachment of a pilus-less bacterium to a solid surface through polymers out of the cell wall, noted as “surface polymers”.

A single, non-specific molecular interaction would unlikely be strong enough to withstand the applied field. Plus, if tethered by a single bond, its breakage would immediately set the cell free, leaving very low probability for repeated attachment. Although unable to visualize these polymers, one can envision multiple tethers with several transient contacts, which best account for the large variation of the drag speed, while the cell remains continually attached. Additionally, since *C. crescentus* has a curved shape, as the cell is frequently going through moments of attachment and detachment, the region of cell-surface contact may vary significantly. The strength of interaction, including that attributable to the DLVO theory, is expected to vary; likewise is the strength of attachment. The bonding formed when different areas of the cell closely interact with the plastic surface could lead to variations in the speed of motion of a crescent-shaped cell even if there is a constant force driving it to move parallel to the surface. Taken together, these considerations may account for the noisy velocity profiles we measured.

## Conclusion

We report on the observation of frequent and transient attachments and detachments of pilus-less mutants of *C. crescentus* from a solid surface. The events were identified by applying an electric field parallel to the surface. These transient attachments and the dragged motion as the cells are repeatedly detached manifest a complex interaction between the bacteria and the solid surface. Our findings point to a weak mode of bacteria-surface interaction, which we refer as “quasi-attachment”. The physical origin of the interaction may be caused by multiple “polymeric tethers”, with individually transient but repeated interactions. The quasi-attachment may be complemented by the non-specific cell body-surface interaction attributable to the classical DLVO theory, but we found qualitatively different behavior from bead-surface interactions presumably lacking polymeric tethers. The model we propose based on the analysis of the “quasi-attachment” events offers mechanistic insights, which guide further experiments, as well as potential applications requiring bacterial attachment to or removal from surfaces.

## Methods

Two strains of *Caulobacter crescentus* were used in this study: CB15 ΔPilin (provided by Yves Brun when he was at Indiana University) and SB3860, which is a mutant derived from CB15 ΔPilin (kindly generated by Bert Ely from University of South Carolina, Columbia). Both bacterial strains lack pilus, making them less likely to adhere to surfaces. SB3860 swims only forward.

*Caulobacter crescentus* was grown in PYE (peptone yeast extract) medium, followed by a synchronization method^25^, in order to select swarmer cells. The culture containing primarily swarmer cells was diluted in DI water, in a proportion of 1:3 in PYE:water, and the mixture was inserted into a microscope slide containing a capillary channel (*μ*-slide I luer from *Ibidi*, Munich, Germany), as illustrated in Figure 7. The channel dimensions are: 50 mm in length, 5 mm in width, and 0.4 mm in height.

**Figure 7.**
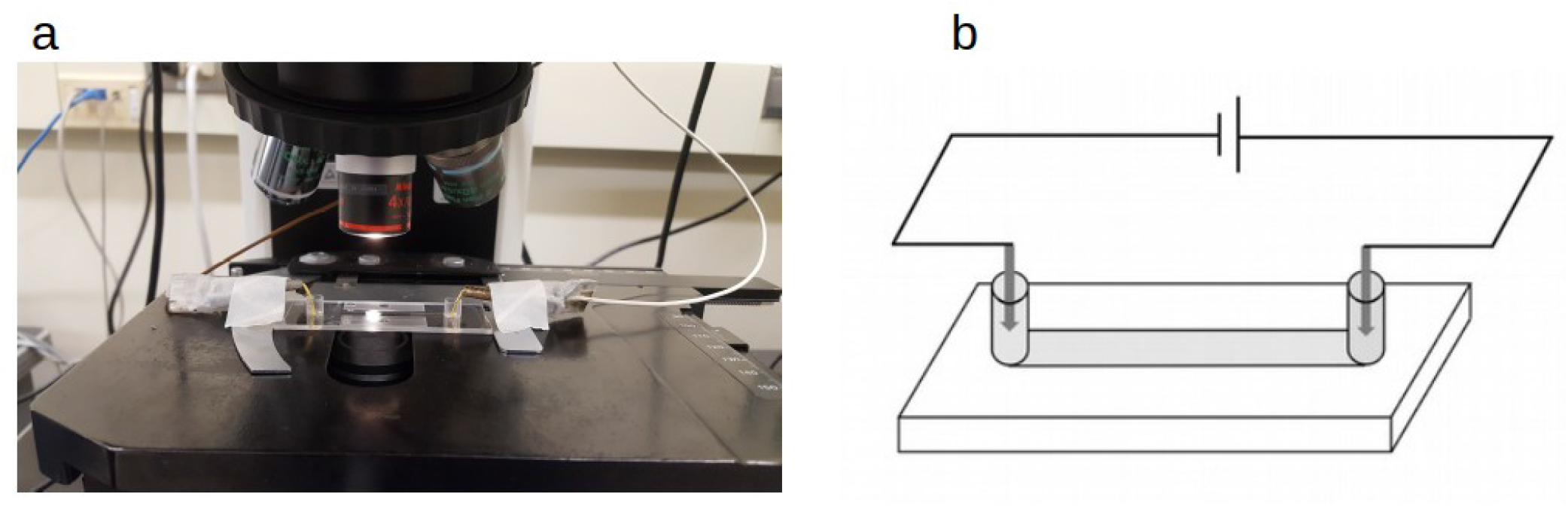
Experimental setup. a) Picture showing a sample slide on a microscope stage. Two wires are connected to electrodes placed at both ends of a capillary filled with the bacteria containing liquid. The microscope slide is fixed so that a part of its capillary channel is imaged while an electric voltage is applied. b) A simple sketch illustrating the electric circuit.

Concentrated suspension of 2.0*μ*m diameter polystyrene beads in 1% volume fraction (Polysciences Inc, Warrington, PA), was first diluted 1000x in DI water, and then to a 1:3 ratio of PYE:water mixture so that in control experiment the beads were subjected to the same ionic condition as the bacteria.

Both ends of the channel were sealed with agarose gel (0.5 % in weight of agarose in a 50 mM *Na_2_SO_4_* solution). Specifically, agarose mix in liquid state was gently deposited at one end of the channel previously filled with the bacteria containing liquid. The slide was gently tilted to aid the gel mix to enter the channel by a millimeter or two. After a 2 minutes wait for the gel to solidify, another drop of agarose mix was filled to the other end of the channel, resulting in both ends of the capillary to be sealed by the agarose gel (see Figure 7 for channel visualization). The agarose gel prevents electrochemical products at the electrodes from spreading into the channel, adversely affecting the bacterial behavior. The agarose seal on both ends of the capillary also efficiently inhibits the fluid flow within the capillary. In addition, the surface of the type of capillaries we purchased is hydrophobic, which effectively suppresses electro-osmotic flow. As a result, we noted no obvious flow near the surface when the electric field was applied during the course of our experiment.

The electrodes were made of gold (99. 99% purity, from Sigma-Aldrich), which is an excellent conductor. This concern of sealing the channels with agarose gel and choosing an inert metal came from the observation that the products of chemical reactions happening at the electrodes were harmful to bacteria and a less restrictive setup had the potential to kill most bacteria within a few seconds under applied voltages over 50 V. DC voltages up to 120 V were provided by a power supply (CSI12001X, Circuit Specialists, Tempe, AZ). We noticed that applying voltages over several minutes invariably caused gas bubbles and even cracks of the agarose gel near the electrodes, where chemical reactions occur. The bubbles produced in these regions progressively altered the electric current. To avoid this adverse effect, most of our experimental data were acquired within a few mins following an applied voltage.

Electric currents were measured by adding an amp-meter (DROK, China) in series to the circuit. Upon application of an electric field, the capillary channel behaved as an Ohmic conductor. We noted, however, that the measured current increased over time, following an applied voltage. In order to obtain proper I-V curve, we averaged measurements by ramping the voltage up and down sequentially in order to eliminate the effect of drift on the shape of the curve. The results the I-V measurements are shown in the supplemental material, along with more detailed description.

Images were acquired using phase contrast under an upright microscope (Eclipse E800; Nikon) with 4x or 40X objective lenses (Nikon), dependent on the size of the region interested. Images were recorded with a charge-coupled device camera (Coolsnap EZ; Photometrics). The acquisition process was controlled using Metamorph software (Molecular Devices). The frame rates for video recordings were either 34 or 10 frames per second (fps).

For image processing, bacteria were tracked using ImageJ and AItracker, an artificial intelligence particle tracker based on neural networks^42^. The bacterial trajectories which were used in the velocity calculations shown in Figures 3 and 4 were processed using a third order Savitzky-Golay filter with window length of 5. This step allows the experimental data to be smoothed, filtering some noise, but keeping its essential features. The tracking of beads was done using the MTrackJ plugin in ImageJ. To smooth out the noise in the calculated velocity of the beads, the running average over 5 consecutive intervals was plotted in Figure 5 for the motion of beads.

## Acknowledgements

We thank Weijie Chen for valuable suggestions. We are grateful to the AItracker (www.aitracker.net) team for processing part of the image tracking data. This work was funded by the National Science Foundation [DMR 1505878]. We also acknowledge financial support from Capes Foundation (Brazil) [graduate fellowship to GA].

## Author contributions

GA and JXT designed the project. GA, JZ and JJO performed the experiments and analysis. GA and JXT wrote the paper.

## Additional information

See Supplementary Information, including the measured I-V curve, average electrophoretic velocities of non-swimming cells measured at 3 applied voltages, an estimation of force applied by the electric field, and the comparison of that force with one produced by flow in a previous study. Supplementary movies 1 and 2 are available in the online version of this paper.

## References

1. Petrova, O. E. & Sauer, K. Sticky situations: Key components that control bacterial surface attachment. J. Bacteriol. 194, 2413–2425 (2012).

2. Dang, H. & Lovell, C. R. Microbial surface colonization and biofilm development in marine environments. J. Bacteriol. 80, 91–138 (2016).

3. Berne, C., Ellison, C. K., Ducret, A. & Brun, Y. V. Bacterial adhesion at the single-cell level. Nat. Rev. Microbiol. 16, 616–627 (2018).

4. Ottemann, K. & Miller, J. F. Roles for motility in bacterial-host interactions. Mol Microbiol 24, 1109–1117 (1997).

5. Donlan, R. M. Biofilms: Microbial life on surfaces. Emerg. Infect. Dis. 8, 881–890 (2002).

6. Jr., W. M. D. Bacterial adhesion: Seen any good biofilms lately? CLINICAL MICROBIOLOGY REVIEWS 15, 155–166 (2002).

7. Rodesneya, C. A., Romana, B., Dhamania, N., Cooleya, B. J., Katirab, P., Touhamic, A. & Gordon, V. D. Mechanosensing of shear by *Pseudomonas aeruginosa* leads to increased levels of the cyclic-di-gmp signal initiating biofilm development. PNAS 114, 5906–5911 (2017).

8. Li, G., Tam, L. & Tang, J. X. Amplified effect of brownian motion in bacterial near-surface swimming. PNAS 105, 18355–18359 (2008).

9. Li, G. & Tang, J. X. Accumulation of microswimmers near a surface mediated by collision and rotational brownian motion. Phys. Rev. Lett 103, 078101 (2009).

10. Li, G., Bensson, J., Nisimova, L., Munger, D., Mahautmr, P., Tang, J. X., Maxey, M. R. & Brun, Y. V. Accumulation of swimming bacteria near a solid surface. Phys. Rev. E 84, 0419327 (2011).

11. Berke, A. P., Turner, L., Berg, H. C. & Lauga, E. Hydrodynamic attraction of swimming microorganisms by surfaces. Phys. Rev. Lett. 101, 038102 (2008).

12. Drescher, K., Dunkel, J., Cisneros, L. H., Ganguly, S. & Goldstein, R. E. Fluid dynamics and noise in bacterial cell-cell and cell-surface scattering. PNAS 108, 10940–10945 (2011).

13. Anderson, B. N., Ding, A. M., Nilsson, L. M., Kusuma, K., Tchesnokova, V., Vogel, V., Sokurenko, E. V. & Thomas, W. E. Weak rolling adhesion enhances bacterial surface colonization. J. Bacteriol. 189, 1794–1802 (2007).

14. Sjollema, J., van der Mei, H. C., Hall, C. L., Peterson, B. W., de Vries, J., Song, L., de Jong, E. D., Busscher, H. J. & Swartjes, J. J. T. M. Detachment and successive re-attachment of multiple, reversibly-binding tethers result in irreversible bacterial adhesion to surfaces. Sci. Reports 7, 4369 (2017).

15. Ipiña, E. P., Otte, S., Pontier-Bres, R., Czerucka, D. & Peruani, F. Bacteria display optimal transport near surfaces. Nat. Phys. 15, 610–615 (2019).

16. Gristina, A. G. Biomaterial-centered infection: microbial adhesion versus tissue integration. Science 25, 1588–1595 (1987).

17. Gómes-Suárez, C., Busscher, H. J. & der Mei, H. C. V. Analysis of bacterial detachment from substratum surfaces by the passage of air-liquid interfaces. App. Env. Microbiol. 67, 2531–2537 (2001).

18. Hong, S. H., Jeong, J., Shim, S., Kang, H., Kwon, S., Ahn, K. H. & Yoon, J. Effect of electric currents on bacterial detachment and inactivation. Biotech. Bioeng. 100, 379–386 (2008).

19. Gall, I., Herzberg, M. & Oren, Y. The effect of electric fields on bacterial attachment to conductive surfaces. Soft Matter 9, 2443–2452 (2013).

20. Poortinga, A. T., Smit, J., van der Mei, H. C. & Busscher, H. J. Electric field induced desorption of bacteria from a conditioning film covered substratum. Biotech. Bioeng. 76, 395–399 (2001).

21. Poindexter, J. S. The caulobacters: Ubiquitous unusual bacteria. MicroBiol. Rev. 45, 123–179 (1981).

22. Skerker, J. M. & Laub, M. T. Cell-cycle progression and the generation of asymmetry in *Caulobacter crescentus*. Nat. Rev. Microbiol. 2, 325–337 (2004).

23. Tsang, P. H., Li, G., Brun, Y. V., Freund, L. B. & Tang, J. X. Adhesion of single bacterial cells in the micronewton range. PNAS 103, 5764–5768 (2006).

24. Bodenmiller, D., Toh, E. & Brun, Y. V. Development of surface adhesion in *Caulobacter crescentus*. J. Bacteriol. 186, 438–1447 (2004).

25. Li, G., Brown, P. J., Tang, J. X., Xu, J., Quardokus, E. M., Fuqua, C. & Brun, Y. V. Surface contact stimulates the just-in-time deployment of bacterial adhesins. Mol. Microbiol. 83, 41–51 (2012).

26. Lele, P. P., Roland, T., Shrivastava, A., Chen, Y. & Berg, H. C. The flagellar motor of *Caulobacter crescentus* generates more torque when a cell swims backward. Nat. Phys. 12, 175–178 (2015).

27. Persat, A., Stone, H. A. & Gitai, Z. The curved shape of *Caulobacter crescentus* enhances surface colonization in flow. Nat. Comm. 5, 3824 (2014).

28. Liu, B., Gulino, M., Morse, M., Tang, J. X., Powers, T. R. & Breuer, K. S. Helical motion of the cell body enhances *Caulobacter crescentus* motility. PNAS 111, 11252–11256 (2014).

29. Morse, M., Bell, J., Li, G. & Tang, J. X. Flagellar motor switching in *Caulobacter crescentus* obeys first passage time statistics. Phys. Rev. Lett. 115, 198103 (2015).

30. Son, K., Guasto, J. S. & Stocker, R. Bacteria can exploit a flagellar buckling instability to change direction. Nat. Phys. 9, 494–498 (2013).

31. McClaine, J. W. & Ford, R. M. Reversal of flagellar rotation is important in initial attachment of *Escherichia coli* to glass in a dynamic system with high-and low-ionic-strength buffers. Appl. Environ. Microbiol. 68, 1280–1289 (2002).

32. Shi, W., Stocker, B. A. & Adler, J. Effect of the surface composition of motile *Escherichia coli* and motile salmonella species on the direction of galvanotaxis. J. Bacteriol. 178, 1113–1119 (1996).

33. Gonin, M., Quardokus, E. M., O’Donnol, D., Maddock, J. & Brun, Y. V. Regulation of stalk elongation by phosphate in *Caulobacter crescentus*. J. Bacteriol. 182, 337–347 (2000).

34. Koyasu, S. & Shirakihara, Y. *Caulobacter crescentus* flagellar filament has a right-handed helical form. J. Mol. Biol. 173, 125–130 (1984).

35. Sharma, S. & Conrad, J. C. Attachment from flow of *Escherichia coli* bacteria onto silanized glass substrates. Langmuir 30, 11147–11155 (2014).

36. Chattopadhyay, S., Moldovan, R., Yeung, C. & Wu, X. L. Swimming efficiency of bacterium *Escherichia coli*. PNAS 103, 13712–13717 (2006).

37. Hormeño, S. & Arias-Gonzalez, J. R. Exploring mechanochemical processes in the cell with optical tweezers. Biol. cell 98, 679–695 (2006).

38. Li, G., Smith, C., Brun, Y. & Tang, J. X. The elastic properties of the *Caulobacter crescentus* adhesive holdfast are dependent on oligomers of n-acetylglucosamine. J. Bacteriol. 187, 257–265 (2005).

39. Walker, S. L., Redman, J. A. & Elimelech, M. Role of cell surface lipopolysaccharides in *Escherichia coli* k12 adhesion and transport. Langmuir 20, 7736–7746 (2004).

40. Li, G. & Tang, J. X. Low flagellar motor torque and high swimming efficiency of *Caulobacter crescentus* swarmer cells. Biophys. J. 91, 2726–2734 (2006).

41. Tsuneda, S., Aikawa, H., Hayashi, H., Yuasa, A. & Hirata, A. Extracellular polymeric substances responsible for bacterial adhesion onto solid surface. FEMS Microbiol. Lett. 223, 287–292 (2003).

42. Newby, J., Schaefer, A., P Lee, M. F. & Lai, S. Deep neural networks automate detection for tracking of submicron scale particles in 2d and 3d. PNAS 115, 9026–9031 (2018).

